# Dynamic Interaction Learning and Multimodal Representation for Drug Response Prediction

**DOI:** 10.1101/2022.11.23.517777

**Authors:** Yanguang Bi, Mu Zhou, Zhiqiang Hu, Shaoting Zhang, Guofeng Lyu

## Abstract

Mining multimodal pharmaceutical data is crucial for in-silico drug candidate screening and discovery. A daunting challenge of integrating multimodal data is to enable dynamic feature modeling generalizable for real-world applications. Unlike conventional approaches using a simple concatenation with fixed parameters, in this paper, we develop a dynamic interaction learning network to adaptively integrate drug and different reactants on multimodal tasks towards robust drug response prediction. The primary objective of dynamic learning falls into two key aspects: at micro-level, we aim to dynamically search specific relational patterns on the whole reactant range for each drug-reactant pair; at macro-level, drug features can be used to adaptively correlate with different reactants. Extensive experiments demonstrate the validity of our approach in both drug protein interaction (DPI) and cancer drug response (CDR) tasks. Our approach achieves superior performance on both DPI (AUC = 0.967) and CDR (AUC = 0.932) tasks, outperforming competitive baselines from four real-world, drug-outcome datasets. In addition, the performance on the challenging blind subsets is remarkably improved, where AUC value increases from 0.843 to 0.937 on blind protein set of DPI task, and Pearson’s correlation value increases from 0.516 to 0.566 on blind drug set of CDR task. A series of case studies highlight the potential generalization and interpretability of dynamic learning in the in-silico drug response assessment.

## I. Introduction

The surge of artificial intelligence (AI) and pharmacology has opened perspectives to drug discovery involving integrative analysis of chemical compound, protein interactions, and gene expression profiles [1]–[4]. For instance, drug protein interaction (DPI) and cancer drug response (CDR) are the two representative tasks of pharmaceutical significance. DPI screens effective chemical compound candidates that have a biological reaction to the specific protein target [3]. Meanwhile, CDR measures patient response to drug candidates based on their biological characteristics [5]. Fundamental questions rise from these scenes as how can we predict the response of drug between various reactants with multimodal learning ability? How do we measure underlying feature interactions to characterize rich drug-target biological connections? Growing efforts have been made using machine learning approaches [3], [6] –[15], yet effective integrative analysis involving molecular, protein, and drug chemical information remains to be addressed explicitly.

Extensive studies have explored individual feature characterization from multi-sourced pharmaceutical data [10], [11], [16]. For instance, drugs are typically understood as a series of chemical formulae, which can be represented as expert-designed molecular fingerprint [9], [16], [17], sentence-style feature [10], [11], and molecular graph structure [3], [6], [7], [12], [18], [19]. In addition, protein sequences and their relational structures, as important sources of drug analysis, have been measured by recurrent neural networks (RNN) [3], [20] and convolutional neural networks (CNN) [9], [10], [12]. To gain insight into integrative power of these quantitative features, it is intuitive to perform feature concatenation. DeepCDR [1] exemplifies the process for concatenating drug feature extraction, cell line feature, and genomics profiles via final fully connected layers. Similar ensemble strategies have been proposed to jointly analyze drug molecular signatures and cell line information [13], [14], [21]. However, such static and rigid feature concatenation with fixed parameters is unable to capture the complex bidirectional interactions to fully reflect pairwise, drug-target responses. Further, the generalization power of modeling various instance drug-target pairs applicable for different tasks of drug prediction is yet to be elucidated.

In this study, we propose the Dynamic Interaction Learning Network (DILNet) to integrate the multimodal features of drug and various reactant towards enhanced response prediction. A key contribution is the dynamic interaction learning module that consists of cascaded depth-wise cross correlations, allowing adaptive interaction patterns extraction for each drug-reactant pair. Such appealing function of DILNet could dynamically refine the interaction parameters for different input pairs. For instance, DILNet can learn the complementary information of key-lock association on different sites in drug-protein interactions [22]. The primary objective of dynamic learning falls into two key aspects: at micro-level, we aim to dynamically search specific relational patterns on the whole reactant range for each drug-reactant pair; at macrolevel, drug features can be used to adaptively correlate with different reactants rather than the choice of fixed and static parameters. The appealing capability of dynamic interaction expands the scope of feature fusion towards generalizable drug-prediction performance. Experimental results highlight that DILNet achieves superior performance on four real-world, drug-outcome datasets, outperforming competitive baselines on both DPI and CDR tasks. In particular, the challenging blind testing results are remarkably improved compared with state-of-the-art baselines, suggesting the potential significance in pharmaceutical applications (see Fig. 3 and Table III). We also have presented two crystal cases in DPI task, where the majority of the binding sites exhibit distinctive values on corresponding feature maps (see Fig. 5). Three representative case studies are further analyzed in CDR task where associated genes in cell lines for specific drugs exhibit strong gradient influence on prediction (Table V). Together, the proposed DILNet demonstrates a dynamics learning of pair-wise interactions that has redefined the search space of drug-reactant features, achieving high-level performance in the context of multimodal DPI and CDR drug-outcome assessment.

**TABLE I.** The quantitative results (mean±standard deviation) of our method and baselines on Human and C.elegans datasets. The best scores are highlighted in bold.

| Datasets | Human |  |  | C.elegans |  |  |
| --- | --- | --- | --- | --- | --- | --- |
| Method | AUC | Recall | Precision | AUC | Recall | Precision |
| kNN | 0.860 | 0.927 | 0.798 | 0.858 | 0.827 | 0.801 |
| RF | 0.940 | 0.897 | 0.861 | 0.902 | 0.844 | 0.821 |
| L2 | 0.911 | 0.913 | 0.891 | 0.892 | 0.877 | 0.890 |
| GNN | 0.970 | 0.918 | 0.923 | 0.978 | 0.929 | 0.938 |
| DrugVQA | 0.979±0.003 | 0.961±0.002 | 0.954±0.003 | - | - | - |
| Our | **0.990±0.002** | **0.962±0.002** | **0.962±0.003** | **0.985±0.003** | **0.963±0.002** | **0.963±0.002** |

## II. Methods

### A. Problem Formulation

Given a pair of drug-target as input, the prediction task is defined to output a drug response result. For the DPI task, the targets are specified as molecular-level proteins. The response output is *y_p_* ∈ {0,1} that denotes a negative or positive interaction between the drug and protein. Therefore, DPI task is formulated as a binary classification problem. For the CDR task, the targets are specified as cancer cell-line data. The output is *y_c_* ∈ ℝ^1^ and commonly measured by the IC-50 value (half-maximal inhibitory concentrations). Therefore, CDR task is formulated as a regression problem in our study.

Generally, drugs are given by the SMILES [23] (Simplified Molecular Input Line Entry System) string. Based on the SMILES syntax, we can restore the molecular graph 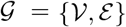 of a drug using RDKit [24], where 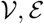 denote the sets of atoms and chemical bonds, respectively. Assuming that 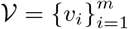 which has *m* atoms with *x*_0_ dimensional descriptor each and 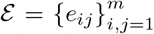 where *e_ij_* denotes the chemical bond between the *i*th and *j*th atoms, we can obtain the graphbased mathematical representations of a drug: feature matrix 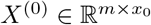 and adjacency matrix *A* ∈ ℝ^*m×m*^.

For the DPI task, the proteins are composed of amino acids, and thus a protein is usually given by the amino acid sequence 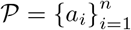 where there are *n* amino acids and *α_i_* denotes the *i*th one. Assuming that each amino acid has *p*_0_ dimensional descriptor, we can obtain the mathematical representation of a protein: feature matrix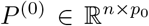. For the CDR task, the cell lines are already structural feature vectors. Recent released dataset [1] has adapted multi-omics profiles to describe cell line such as **g**enomic, **t**ranscriptomic, and **e**pigenomic. Assuming that the dimensions of above features are *g, t, e*, we can obtain the mathematical representations of a cell line: feature matrices *C_g_* ∈ ℝ^1×*g*^, *C_t_* ∈ ℝ^1×*t*^, and *C_e_* ∈ ℝ^1×*e*^.

### B. Dynamic Interaction Learning Network

Fig. 1 illustrates core components of the proposed DILNet for drug response prediction. The upper left (blue box) is the drug graph feature module and the lower left (green box) is the target sequence feature module. These initial extracted features are integrated by the dynamic interaction learning module (yellow box) for the final prediction task (cyan box). To extend the scope of the framework, the target sequence feature extraction (green box) can be replaced by the cancer cell line inputs for the CDR task, while the integration learning module remains unchanged.

**Fig. 1.**
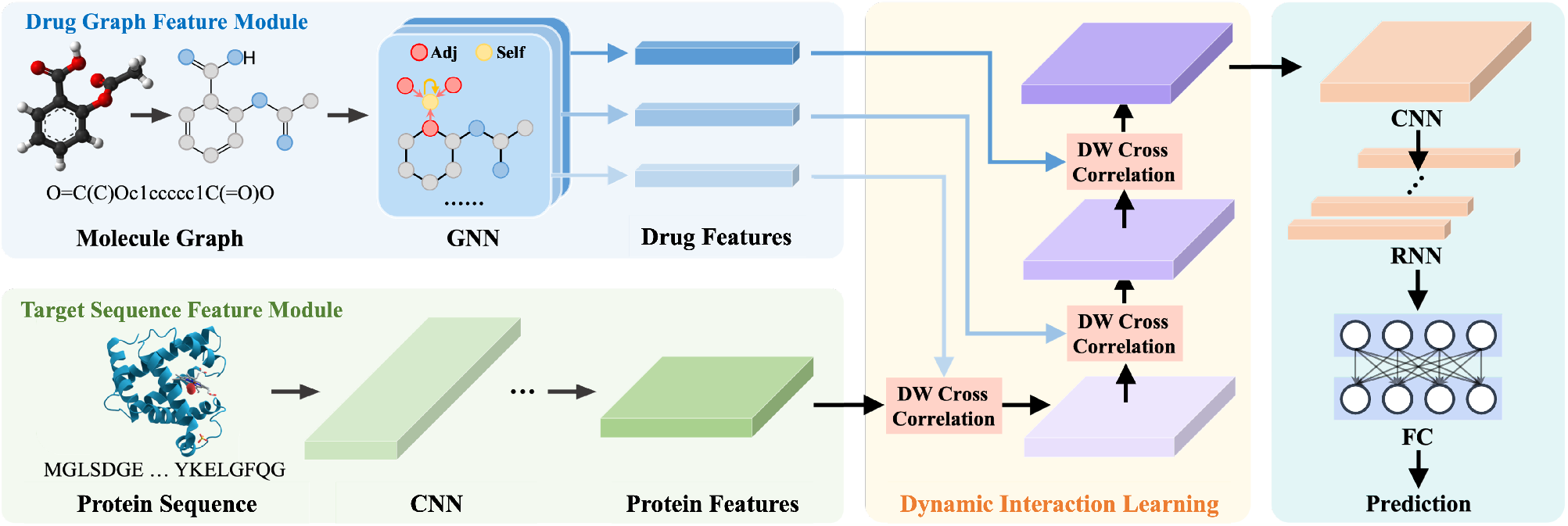
The overview of the framework for the DPI task. Individual representations for the drug (blue box) and the protein (green box) are fused by the dynamic interaction learning module (yellow box) for the following prediction (cyan box). For other tasks, the protein representation learning module (green box) can be readily replaced to fit the task-specific target, while the main dynamic interaction learning module remains the unchanged. Next, the interaction learning module performs DW cross correlation to enable feature integration, and prediction module accomplishes drug outcome response prediction.

DILNet learns individual features for each drug-target pair respectively. For the drug 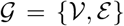 where 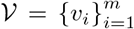, denote the descriptors for the *m* atoms as 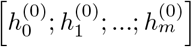. In this study, the drug descriptors are the extracted 75-dimensional vectors [25], including both chemical and topological properties such as atom type, degree, and hybridization. To extract drug graph-level molecular features, we use Graph Isomorphism Network (GIN) [26] due to its proven discriminative power. Formally, GIN updates the node features as

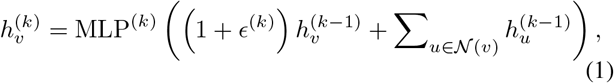

where 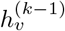 denotes hidden features of node *v* in the (*k* – 1)th layer, 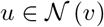 denotes the neighbor node of v determined by the adjacency matrix. *ϵ* is a learnable parameter to control the self weight of current node *v*. The aggregation is followed by a Multi-Layer Perceptron (MLP) to produce the hidden features 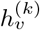 for the kth layer. The updates proceed iteratively, where the intermediate features at different depth are applied average pooling over atoms to get the group of final features {*X*^(*k*)^},

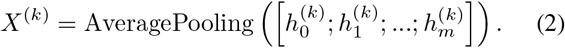

For the protein 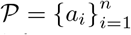, assuming that each amino acid has *p* dimensional descriptors, the protein can be collectively represented as *P*^(0)^ ∈ ℝ^*n×p*^. In this work, we simply take the one-hot representation of amino acids as the descriptors. Overall, there are 23 types of amino acids (20 standards, 2 additional, 1 for unknown). For instance, taking *P*^(0)^ ∈ ℝ^*n×p*^ as a sequence with *n* elements and *p* channels, we design a simple 1D-CNN in target sequence feature module to extract the features. The CNN contains four stages where each stage includes three groups of multi-scale convolution layers (MS Convs) and a pooling layer (removed for the last stage), as shown in Fig. 2. The design is inspired by [20], [27] to adaptively capture the sequence features at different distances. The output feature of the last stage is taken as the protein feature and denoted as *P*.

**Fig. 2.**
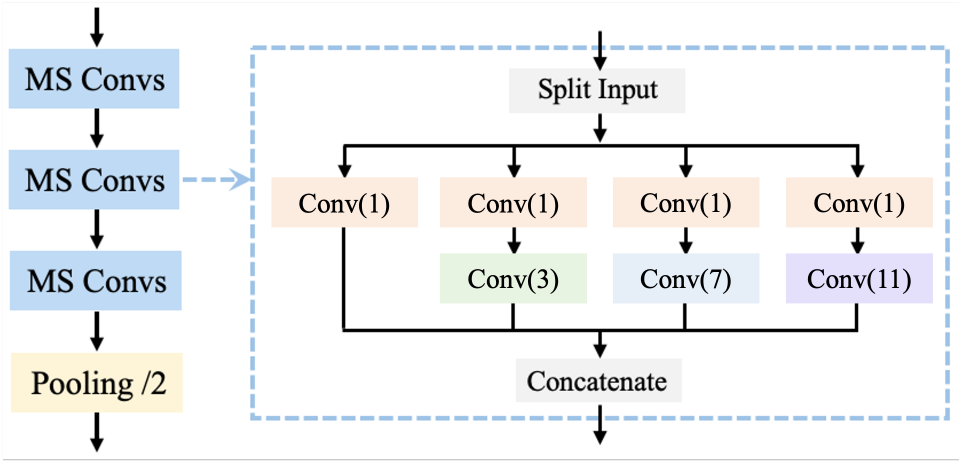
Illustration of the convolutional operations of our 1D-CNN architecture. The convolutional pipeline contains three groups of multi-scale convolution layers (MS Convs) and a pooling layer. The “Conv(n)” includes the batch normalization and rectified linear unit, and *n* denotes kernel size.

To seek the generalization power of DILNet, we perform module design to be applicable for the second task of CDR analysis. As seen in Fig. 1, for the target sequence feature module, we only use the genomic, transcriptomic, and epigenomic features defined as 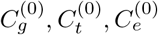, respectively to replace the protein input in DPI task. These features are firstly mapped to the fixed dimension *d*, and concatenated as *C*^(0)^ ∈ ℝ^*d*×3^. *C*^(0)^ is seen as a sequence with d elements and 3 channels, and we apply an 1D-CNN with three convolution layers and two pooling layers. The output feature is thus taken as the cell line feature and denoted as *C*.

Given individual features {*X*^(*k*)^} produced by drug graph feature module and *P* produced by the target sequence feature module, the dynamic interaction learning module (yellow box in Fig. 1) aggregates the features to give the joint representation. Notably, the architecture of the interaction learning module is consists of cascaded depth-wise cross correlations (DWCCs) without parameters. Assuming that DWCC has the input *z* ∈ ℝ^*z×c*^ and the kernel *x* ∈ ℝ^*x×c*^ where *z,x* and *c* denote the input size, kernel size and number of channels, the output *s* = DWCC (*x, z*) is given by

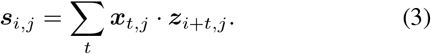

DWCCs are stacked to model the complex bilateral interactions, and can be expressed as

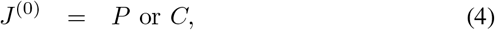

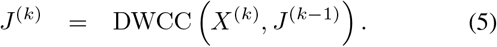

Note that kernels in the hierarchical DWCCs come from drug features {*X*^(*k*)^} outputted by GNN at different depth *k*. As a result, these features aggregate information from different neighborhood ranges within the graph. This is analogy to the multi-scale learning in CNNs, and characterizes bilateral interactions at different granularities.

Finally, the dynamically integrated feature *J*^(*K*)^ produced by the interaction learning module with *K* iterations is analyzed by the prediction module to make the final prediction. For the DPI task, the prediction module firstly employs another 1D-CNN *ϕ_c_* to further fuse the tracking features as the transition for next RNN *φ_r_* which extracts the hidden interaction states. Then, FC layers *φ_f_* followed by Sigmoid function *σ* are used to obtain the final classification score *Y* ∈ ℝ. The entire prediction module is a cascaded structure as follows:

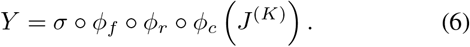

For the CDR task, we remove the RNN *ϕ_r_* and Sigmoid function *σ*, and the structure can be summarized as,

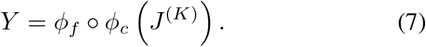

Overall, the DILNet is end-to-end trainable and supervised by binary cross entropy loss for DPI classification task or RMSE (root mean square error) loss for CDR regression task.

## III. Experimental results

### A. Datasets

#### Human

dataset [2] contains 3,369 positive interactions between 1,052 unique compounds and 852 unique proteins. **C.elegans** dataset contains 4,000 positive interactions between 1,434 unique compounds and 2,504 unique proteins. Following the convention, the balanced datasets [11], [12] are used where positive and negative interactions are 1:1 and randomly split 80%, 10% and 10% for training, validation, and testing.

#### BindingDB

[28] is a public, web-accessible database that contains massive real-world measured binding affinities between small molecule drugs and proteins. We use the customized BindingDB dataset [3] which takes a snapshot from the original entire BindingDB database. There are 39,747 positive interactions and 31,218 negative interactions which are divided into three subsets: training set (50,155 interactions), validation set (5,607 interactions) and testing set (5,508 interactions). Note that about 45% proteins in the testing set are not seen in the training set.

#### DeepCDR

[1] is a recently released public dataset for the CDR task, which aggregates multi-omics profiles from GDSC (Genomics of Drug Sensitivity in Cancer) [29], CCLE (Cancer Cell Line Encyclopedia) [30], and TCGA (The Cancer Genome Atlas) [31] databases. It contains 107,446 instances across 561 cancer cell lines and 238 drugs, where each instance denotes a pair of drug and cancer cell line with their response IC50 value (natural log-transformed). The 95%/5% of the dataset is split for training/testing as its default setting.

### B. Implementation Details

In the model architecture, all the hidden layers in GIN have 64 output channels. The CNN for protein representation learning has four convolution stages where the output channels are 16, 32,48, and 64. The pooling layer in the last convolution stage is removed. For the second CDR task, the cell line feature learner CNN contains three convolution layers with output channel 128 and two pooling layers. The prediction module is slightly different between the DPI task and the CDR task. For the DPI task, joint features are firstly processed by CNN which has two convolution stages with 128, 256 output channels, and no pooling layers. For the following RNN we employ Bi-LSTM [32] with 128 hidden size, and finally we add three FC layers with 256 hidden size as the classifier. The prediction module for the CDR task has been reformulated, where CNN contains three convolution layers and two pooling layers as similar to the cell line feature learner.

For the training procedure, we use Adam optimizer [33] with 1 × 10^-5^ weight decay setting. The initial learning rate is 1 × 10^-3^ and decreases to 1 × 10^-5^. The model is trained 100 epochs with batch-size 32. Model selection and hyperparameters are determined by the validation results. Further, for the comparison with baseline approaches, we directly refer their reported results in the literature [1], [11], [14], [15].

### C. Benchmark Comparisons

#### Drug Protein Interaction

Table I shows that our approach achieves the leading performance in both Human and C.elegans datasets. The mean and std standard deviation are both reported over 10 repetitions for fair comparison. We have referred results of kNN (k-nearest neighbor), RF (random forest), L2 (L2-logistic), and GNN (graph neural network) from [12] and the result of DrugVQA is from [11]. It is noted that the results of traditional machine learning methods (kNN, RF, L2) are under performed comparing with graph-driven approaches (e.g., GNN and DrugVQA). These graph-driven examples have shown the promise of mining graph structured information, but they adopt the naive feature stacking learning. In contrast, the proposed DILNet explores the latent biochemical interaction mechanism to boost results on all metrics.

We also evaluate our approach in the BindingDB dataset against other state-of-the-art methods, including Tiresias [34], DBN [8], GNN [12], E2E [3], DrugVQA [11]. Following [3] and [11], we present results for “proteins are seen” and “proteins are novel” subsets respectively in the histogram form as seen in Fig. 3(a). For the proteins-seen subset, all the methods attain promising performance (AUC ¿ 0.9) and our method achieves the highest performance with AUC of 0.986. For the proteins-unseen subset, however, performances of Tiresias, DBN, GNN drop sharply. In comparison, our approach maintains a high-level performance with AUC of 0.937, demonstrating the strong prediction ability on capturing drug-protein associations.

**Fig. 3.**
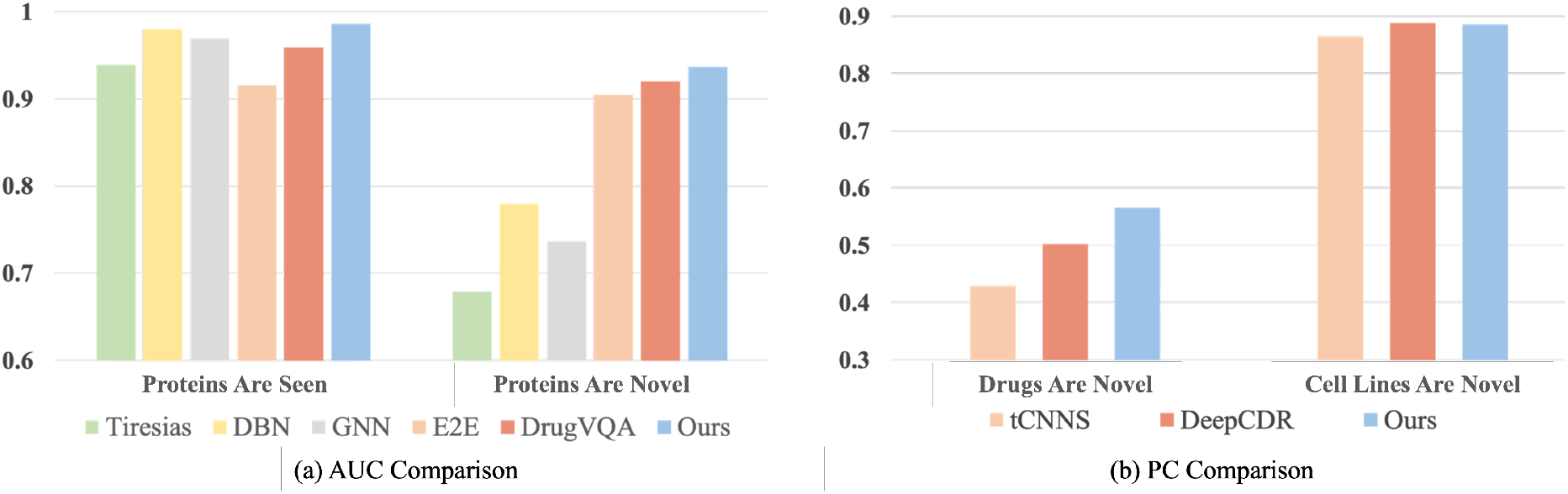
Performance comparison of our method versus other state-of-the-art methods in (a) the BindingDB dataset of the DPI task evaluated by AUC. (b) the DeepCDR dataset of the CDR task evaluated by pearson’s correlation (PC).

#### Cancer Drug Response

We extend to perform CDR predictive analysis on the DeepCDR dataset. As seen in Table II, we compare quantitative results with multiple baselines including Ridge Regression [35], Random Forest [35], MOLI [15], CDRscan [13], tCNNS [14], and DeepCDR [1]. Similar to the DPI task, the mean and std standard deviation of CDR task are also reported over 10 repetitions for fair comparison. Our method achieves higher performance than the SOTA DeepCDR in all metrics including RMSE, pearson’s correlation (PC), and spearman’s correlation (SC). To our best knowledge, the value of RMSE is for the first time lower than 1 compared with prior studies in CDR analysis. Such finding indicates that the proposed DILNet demonstrates potential generalization power to effectively adapt to the CDR task with promising results.

**TABLE II.** The quantitative results (mean±standard deviation) of our method and baselines on DeepCDR dataset. The best scores are highlighted in bold.

| Method | RMSE | Pearson's Correlation | Spearman's Correlation |
| --- | --- | --- | --- |
| Ridge Regression | 2.368 | 0.780 | 0.731 |
| Random Forest | 2.270 | 0.809 | 0.767 |
| MOLI | 2.282±0.008 | 0.813±0.007 | 0.782±0.005 |
| CDRscan | 1.982±0.005 | 0.871±0.004 | 0.852±0.003 |
| tCNNS | 1.782±0.006 | 0.885±0.008 | 0.862±0.006 |
| DeepCDR | 1.058±0.006 | 0.923±0.006 | 0.903±0.004 |
| Our Method | **0.993±0.003** | **0.932±0.004** | **0.907±0.003** |

Fig. 3(b) presents additional blind test comparisons for unseen drugs and unseen cell lines with new dataset splits. The PC in blind cell line test is slightly lower than the baseline (DeepCDR). However, for the much more challenging blind drug test where the baselines are largely depressed, the PC of the proposed method is largely improved to 0.566 compared with the 0.503 of DeepCDR by a notable performance margin.

### D. Ablation Study

#### Dynamic Interaction Learning versus Concatenation Baseline

To eliminate the effects by individual representation learning modules of different methods, we here replace the proposed dynamic integration learning module with feature concatenation as the baseline. Meanwhile, the remained modules and training procedures are kept the exactly unchanged.

Table III displays that the concatenated baseline achieves a highly similar result (AUC 0.933) as to the DeepVQA (AUC 0.936) in the DPI task; and that is also comparable (PC 0.925) with the DeepCDR (PC 0.923) in the CDR task. This finding indicates that difference in the individual representation learning modules does not account for the significant overall performance difference. In addition, when further comparing the feature concatenation baseline with the proposed dynamic integration learning, we identify that the performance improvement is remarkably recognized in novel-reactant scenarios. In the DPI task, two methods are both accurate enough when proteins are seen, but in novel-protein scenario, dynamic integration learning outperforms concatenation baseline with a clear margin (0.843 to 0.937 in AUC and 0.757 to 0.854 in ACC); similarly in the CDR task, performances are comparable for the novel-cell line scenario but dynamic integration learning shows significant advantage in the more challenging novel-drug scenario. These results suggest that dynamic interaction learning is promising towards enhanced prediction of drug outcomes under different settings.

**TABLE III.**
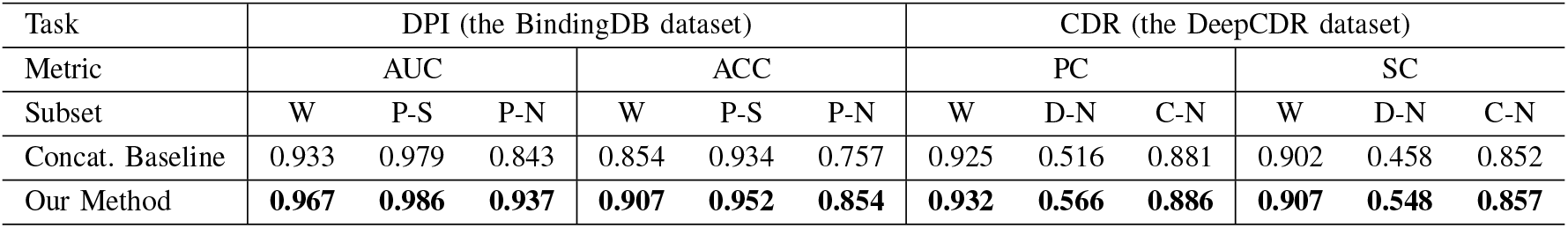
Performance comparison of the proposed dynamic integration learning and the concatenation baseline on different subsets of the DPI and CDR tasks. “W”, “P-S”, “P-N”, “D-N” and “C-N” denote the whole testset, proteins-seen subset, proteins-novel subset, drugs-novel subset, and cell lines-novel subset, respectively. The leading scores are highlighted in bold.

#### Number of Stacking DWCCs

We further investigate how the number of deep-wise cross correlation (DWCCs) will affect the overall performance. We perform experiments in both two tasks based on BindingDB and DeepCDR datasets. The results are shown in Table IV where we find that multiple stacking DWCCs consistently outperform the single DWCC. This is analogous to the superiority of MLP with the increased expressive power over single-layer classifier. Also similar to MLP, the results are not necessarily better with more layers, thus the number of DWCCs should be tuned to fit diverse practical tasks.

**TABLE IV.**
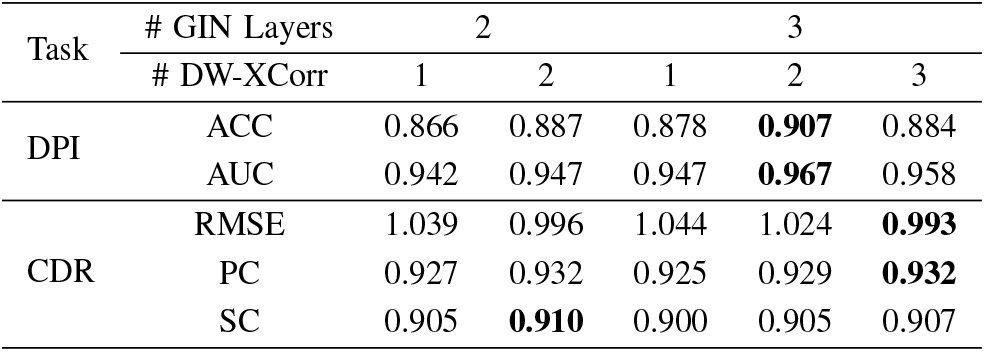
The quantitative results of our method with different configurations in the pattern tracking module for DPI (BindingDB Dataset) and CDR (DeepCDR Dataset) tasks. The best scores are highlighted in bold.

### E. Case Study and Interpretation

We perform case study on both DPI and CDR tasks to further assess model utility of our approach in the context of biochemical interpretation. For the DPI task, the 3OXZ crystal structure [36] and another 3TGG crystal structure [37] are selected from PDB (Protein Data Bank) [38] as illustrative examples. It is known that 3OXZ contains a positive interaction with strong binding affinity between protein “Tyrosine-protein kinase ABL1, UniProtKB-P00520” and drug “Ponatinib, C_29_H_27_F_3_N_6_O”. Our approach produces 0.989 score which correctly predicts the interaction of unseen protein. Similarly, 3TGG contains a positive interaction between protein “Phosphodiesterase, UniProtKB-O76074” and drug “C_23_H_31_N_5_O_4_”, which is also successfully predicted with 0.961 score in our study. During model inference, the intermediate tracking features are additionally extracted and compressed into 1D using principal component analysis (PCA) for visualized observation. The intermediate feature map has 1/8 length of the original amino acid sequence due to pooling layers. As shown in Fig. 5, the 3D binding sites are mapped into the corresponding 1D amino acid sequence where 3/4 binding sites (numbers 1, 3, 4) in 3OXZ and 1/1 binding site in 3TGG exactly have particularly distinguish feature values in their 1D locations. The findings demonstrate that the proposed DILNet is able to dynamically capture the unique interaction patterns between different drug-protein pairs, which contributes to the interpretable prediction with improved performance.

**Fig. 4.**
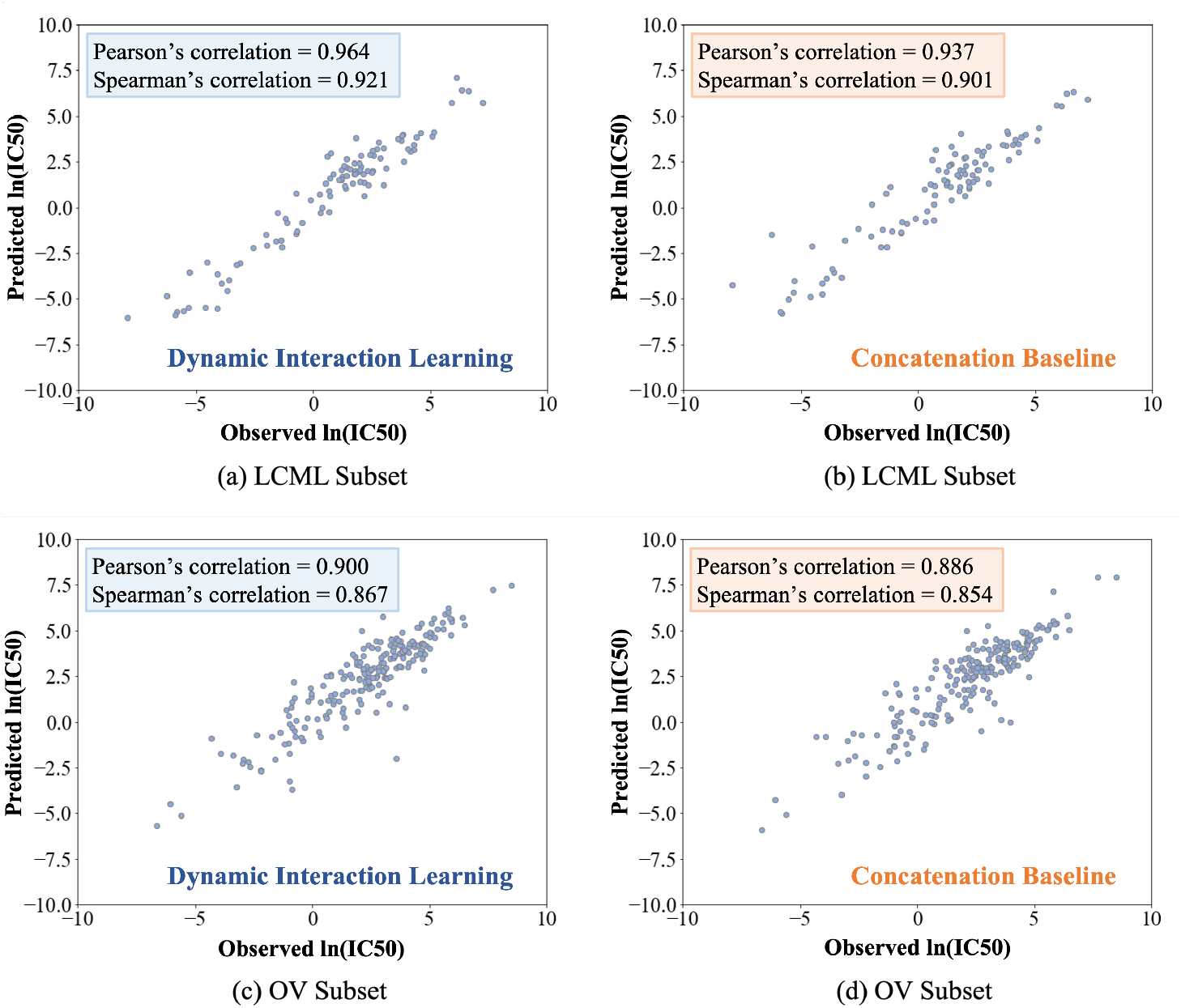
Predicted and Observed IC50 values of our method and concatenation baseline on (a)-(b) LCML subset with the best performance and (c)-(d) OV subset with the worst performance in the DeepCDR dataset.

**Fig. 5.**
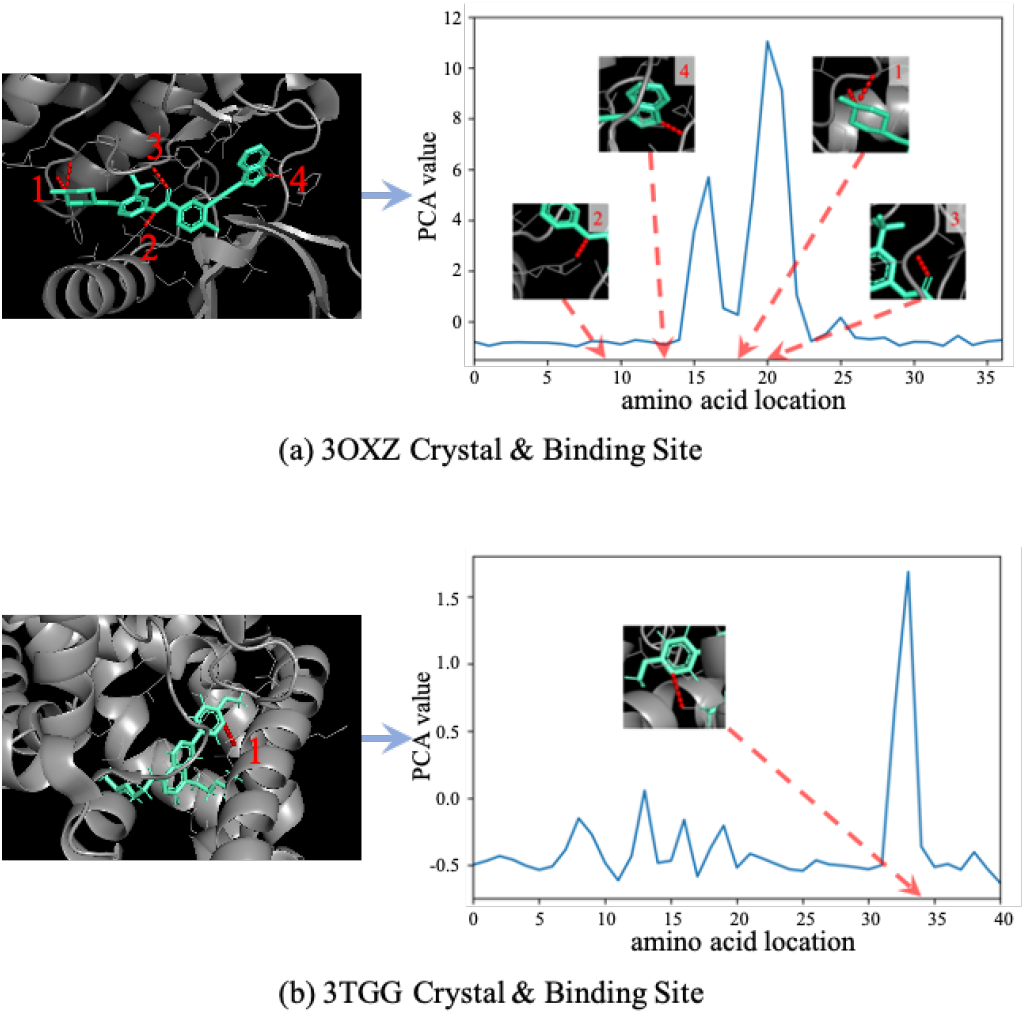
The crystal structures and binding sites in corresponding feature map. (a) 3OXZ crystal structure and 4 binding sites. (b) 3TGG crystal structure and 1 binding site. The red arrows denote 3D ligand sites and their corresponding 1D locations in the amino acid sequence. Overall, the proposed method successfully captures most binding sites for interpretable prediction.

For the CDR task, since the DeepCDR cohort is usually split into different clusters in terms of the TCGA cancer types, we here select the LCML (Chronic Myelogenous Leukemia) cluster subset and the OV (Ovarian Serous Cystadenocarcinoma) cluster subset as the representative cases, which respond to the highest and the lowest results respectively in our study. As visualized in Fig. 4, our method achieves better regression performance compared with the concatenation baseline in both clusters. Table V presents interpretable response prediction between the drug and cell line following the setting in DeepCDR [1]. We found that the observed and predicted IC50 values are strongly aligned, suggesting well generalization ability to measure in-vitro cancer drug response. We also calculate the gradient of each gene (697 totally) with respect to final output score when the drug data is fixed. Then, the output of top-1 gene is ranked based on the modulus of corresponding gradient. For example, the PIM1 gene ranks first in the response between Docetaxel and PF-382. Such finding is supported by the existing literature [39], which reveals that the PIM1 kinase is a critical component of a survival pathway activated by Docetaxel and promotes survival of Docetaxel-treated prostate cancer cells. Similarly, we show that the KLF6 gene ranks first in the model response between Bortezomib and SUP-B15, which is supported by existing literature [40] that Bortezomib significantly upregulated the expression of KLF6. In addition, the PTEN gene also ranks first in the model response between Elesclomol and BFTC-905, while [41] shows that Elesclomol contributes to the treatment of breast cancer cells with a lack of PTEN. Taken together, the unique integration patterns between drugs and associated genes in cell line have been well captured by the proposed DILNet for interpretable response prediction.

**TABLE V.** Top-1 ranked gene in cell line responded to drug, which is predicted by the proposed meta-learning framework using gradient-based strategy.

| Drug | Cell Line | Top-1 Gene | -ln(IC50) |  |
| --- | --- | --- | --- | --- |
|  |  |  | Observed | Predicted |
| Docetaxel | PF-382 | PIM1 | 2.905 | 3.155 |
| Bortezomib | SUP-B15 | KLF6 | 7.403 | 6.126 |
| Elesclomol | BFTC-905 | PTEN | 3.782 | 3.710 |

## IV. Conclusion

Multimodal representation and feature interaction are crucial for developing robust in-silico models for screening and measuring efficiency of drug delivery. We have proposed the end-to-end Dynamic Interaction Learning Network (DILNet) to adaptively model the complex relations between individual drug and targets. DILNet emphasizes the dynamic learning ability to capture relational measurement and subsequent feature integration of drug and target characteristics. As a result, our task-agnostic approach has demonstrated competitive performance on both DPI and CDR tasks using four public real-world datasets. Interpretable case studies strengthen the utility of our approach to capture the deep integration patterns that contribute to the overall prediction. The performance on novel-reactant subsets are further remarkably improved, indicating the potential generalization power in these drug-sensitive scenarios. Given the rapid growth of pharmaceutical data sources, we plan to investigate the broad validity of DILNet involving different drug-associated data such as single cell sequence data, metabolomics, and proteomics data.

